# Marker-assisted selection of trees with *MALE STERALITY 1* in *Cryptomeria japonica* D. Don

**DOI:** 10.1101/2020.05.29.114140

**Authors:** Yoshinari Moriguchi, Saneyoshi Ueno, Yoichi Hasegawa, Takumi Tadama, Masahiro Watanabe, Ryunosuke Saito, Satoko Hirayama, Junji Iwai, Yukinori Konno

**Author notes:** corresponding author Yoshinari Moriguchi.

## Abstract

Practical use of marker-assisted selection (MAS) is limited in conifers because of the difficulty with developing markers due to a rapid decrease in linkage disequilibrium, the limited genomic information available, and the diverse genetic backgrounds among breeding material collections. First, in this study, two families were produced by artificial crossing between two male-sterile trees, Shindai11 and Shindai12, and a plus tree, Suzu-2 (*Ms1/ms1*) (S11-S and S12-S families, respectively). The segregation ratio between male-sterile and male-fertile trees did not deviate significantly from the expected 1:1 ratio in either family. These results clearly suggested that the male-sterile gene of Shindai11 and Shindai12 is *MALE STERALITY 1* (*MS1*). Because some markers reported previously have not been linkage mapped, we constructed a partial linkage map of the region encompassing *MS1* using the S11-S and S12-S families. For the S11-S and S12-S families, 19 and 18 markers were mapped onto the partial linkage maps of *MS1* region, respectively. There was collinearity (conserved gene order) between the two partial linkage maps. Two markers (CJt020762_*ms1-1* and reCj19250_2335) were mapped to the same position as the *MS1* locus on both maps. Of these markers, we used CJt020762 for MAS in this study. According to the MAS results for 650 trees from six prefectures of Japan (603 trees from breeding materials and 47 trees from the Ishinomaki natural population), five trees in Niigata Prefecture and one tree in Yamagata Prefecture had heterozygous *ms1-1*, and three trees in Miyagi Prefecture had heterozygous *ms1-2*. The results obtained in this study suggested that there may be geographical hotspots for the *ms1-1* and *ms1-2* alleles. Because MAS can be used effectively to reduce the labor and time required for selection of trees with a male-sterile gene, the number of breeding materials should increase in the future.

## Introduction

Molecular marker-assisted selection (MAS), which can reduce the time required for a breeding cycle, is an attractive method for conifers, which have longer generation times than those of most crop species [1]. However, in conifers, practical use of MAS is limited because it is difficult to develop markers for MAS due to a rapid decrease in linkage disequilibrium, the limited genomic information available, and the diverse genetic backgrounds among breeding material collections. Nevertheless, the progress with genome analysis technologies has recently accelerated, producing an enormous volume of sequences and subsequent development of markers linked to a particular target gene.

Sugi (*Cryptomeria japonica* D. Don) is an important forestry species that occupies nearly 4.5 million hectares of artificial forest in Japan, which corresponds to approximately 44% of all artificial forest area in the country [2]. The forestry-related increase in the area covered by *C. japonica* has triggered pollinosis. *C. japonica* pollinosis is one of the most serious allergies in Japan, affecting 26.5% of the Japanese population [3]. As a countermeasure against *C. japonica* pollinosis, male-sterile trees can be implemented effectively. The first *C. japonica* tree with genetic male sterility conferred by a major recessive gene, *MALE STERALITY 1* (*MS1*), was found in Toyama Prefecture in 1992 (Toyama-1) [4, 5, 6]. Since the discovery of this individual, six male-sterile trees homozygous for *MS1* (*ms1/ms1*) have been selected (Shindai3, Fukushima1, Fukushima2, Tahara-1, Sosyun, and Miefunen-1) [7, 8, 9, 10, 11, 6, 12]. The frequency of these male-sterile trees in the forest is considered to be very low, because Igarashi et al. [7] identified only two male-sterile trees in a screening of 8,700 trees distributed across a 19-ha artificial forest. Male-sterile trees are generally identified by observing pollen release and/or by direct inspection of the male strobili using a magnifying glass or microscope. In the selected male-sterile trees, confirmation of the male-sterile gene *MS1* was made based on the results of test crossings. These test crossings led to the discovery of three other male-sterile genes: *MS2, MS3*, and *MS4* [5, 13, 14, 6]. In some male-sterile trees such as Shindai11 and Shindai12, male-sterile genes have not yet been investigated.

Mutations in the *MS1* gene leads to the collapse of microspores after separation of pollen tetrads [15], whereas that of the *MS2* gene leads to the formation of microspore clumps after normal microsporogenesis [13]. On the other hand, mutations in the *MS3* and *MS4* genes lead to the formation of microspores of various sizes after normal microsporogenesis [13, 14]. The four male-sterile genes *MS1, MS2, MS3*, and *MS4* have been mapped to different linkage groups: the ninth (referred to as LG9 hereafter), fifth, first, and fourth linkage groups, respectively [15, 16, 17]. Only one tree with *ms2, ms3*, and *ms4* was selected, respectively. Therefore, trees with *ms1* have generally been used for tree improvement and seedling production. Both male-sterile trees and also trees heterozygous for the male-sterile gene are important for tree improvement and seed production as the maternal and paternal parents, respectively. Currently, seven trees heterozygous for *MS1* (*Ms1/ms1*), Suzu-2, Naka-4, Ooi-7, Ohara-13, Zasshunbo, Kamiukena-16, and Kurihara-4, have been selected [4, 6, 19, 20, 12; Konno, personal communication]. For precise selection of trees heterozygous for *MS1*, it is generally necessary to produce F_1_ trees by artificial crossing and to confirm whether these F_1_ trees are male-sterile or -fertile trees. Confirmation is performed by direct inspection of male strobili using a magnifying glass (or a microscope) or by observing pollen release.

Due to the large amount of labor required for selection, the number of trees with the male-sterile gene is not sufficient. To reduce the labor of screening, MAS of trees with the male-sterile gene is necessary. Recently, some markers closely linked to the *MS1* gene or derived from a putative *MS1* gene have been developed [18, 21, 22, 23]. Moriguchi et al. [18] and Ueno et al. [23] reported that estSNP04188 and dDcontig_3995-165 were 1.8 cM and 0.6 cM from *MS1* in the T5 family (173 trees), respectively. Hasegawa et al. [21] reported that 15 markers were 0 cM from *MS1* in the F1O7 family (84 trees). Among these, AX-174127446 showed a high rate of predicting trees with *ms1*. Mishima et al. [22] reported two markers from contig “reCj19250” that can be used to select trees with *ms1*. On the other hand, Hasegawa et al. [24] reported a candidate male-sterile gene CJt020762 at the *MS1* locus, and all breeding materials with the allele *ms1* had either a 4-bp or 30-bp deletion in the gene (they defined these alleles as *ms1-1* and *ms1-2*, respectively). Both of these were expected to result in faulty gene transcription and function; therefore, they developed two markers [30] from contig “CJt020762”. Some of these markers have not been mapped on a linkage map. The lack of a linkage map for these markers constructed from the same family makes it difficult to understand the relative position of each marker.

Therefore, in this study, we (1) checked whether the male-sterile gene of Shindai11 and Shindai12 is *MS1* based on the results of test crossings, (2) constructed a partial linkage map of the region encompassing *MS1*, and (3) selected trees with *ms1* by MAS. As there are few studies pertaining to practicable applications of MAS in conifers, this study should provide a valuable model.

## Materials and Methods

### Phenotyping of male sterility and SNP genotyping for linkage analysis

We used two families, S11-S and S12-S, in this study. These families were produced by artificial crossing between two male-sterile trees, Shindai11 and Shindai12, and a plus tree, Suzu-2 (*Ms1/ms1*), during March of 2016. Strobili production was promoted by spraying the trees with gibberellin-3 (100 ppm) in July 2018. Approximately five male strobili were sampled from each individual from early November to early December 2018. Each sampled male strobilus was bisected vertically with a razor, and male sterility was determined using a microscope (SZ-ST, Olympus, Tokyo, Japan). Individuals without male strobili and individuals in whom it was difficult to discriminate male sterility were excluded from further analysis. Finally, 130 individuals from S11-S and 138 individuals from S12-S were used to construct a linkage map. Needle tissue was collected from three parent trees (Shindai11, Shindai12, and Suzu-2) and all F_1_ trees (268 trees) of two mapping populations. Genomic DNA was extracted from these needles using a modified hexadecyltrimethylammonium bromide (CTAB) method [25, 30].

Single nucleotide polymorphism (SNP) markers from contigs “reCj19250” and “CJt020762” [22, 24] and SNP markers mapped to LG9 [26, 21, 23] were used to construct a partial linkage map of the region encompassing the *MS1* locus for each of the two families (because the gene is located in LG9) [16]. For estSNP00204 [18], AX-174127446 [21], and CJt020762 [24], the SNaPshot assay, which extends primers by a single base, was used for genotyping. The primer sequences used to target the three markers in the SNaPshot assay (estSNP00204 [18], AX-174127446 [21], and CJt020762 [24]) are shown in Table S1. Although CJt020762 contained a 4-bp and 30-bp deletion, we used the 4-bp deletion for primer design because there is no polymorphism associated with the 30-bp deletion between parents of the mapping populations. Multiplex polymerase chain reaction (PCR) was performed using three primer pairs and the Multiplex PCR Kit (QIAGEN, Hilden, Germany). Each reaction contained 2× QIAGEN multiplex PCR master mix, 1 μL primer mix (2.5 μM for each primer), and 40 ng genomic DNA in a total volume of 6 μL. Amplification was performed in the Takara PCR Thermal Cycler (Takara, Tokyo, Japan) using an initial denaturation step at 95 °C for 15 min, followed by 30 cycles of denaturation at 94 °C for 30 s, annealing at 57 °C for 1.5 min, and extension at 72 °C for 1 min, with a final extension at 60 °C for 30 min. To remove any primers and dNTPs, 5.0 μL of the PCR products were treated with 2.0 μL ExoSAP-IT reagent (Thermo Fisher Scientific, Waltham, MA, USA), followed by incubation at 37 °C for 30 min and then 80 °C for 15 min to inactivate the enzyme. Single-base extension reactions were carried out in a 5.0 μL final volume containing 0.5 μL SNaPshot Multiplex Ready Mix (Thermo Fisher Scientific), 1 μL primer mix (1.0 μM for each primer), and 2.0 μL of the treated PCR products. Reactions were performed in the Takara PCR Thermal Cycler (Takara) with 25 cycles of denaturation at 96 °C for 10 s and annealing and elongation at 60 °C for 30 s. The final extension products were treated with 1 U shrimp alkaline phosphatase (Thermo Fisher Scientific) and incubated at 37 °C for 1 h, followed by enzyme inactivation at 80 °C for 15 min. The PCR products (1.0 μL) were mixed with 0.2 μL GeneScan 120 LIZ size standard and 8 μL Hi-Di formamide prior to electrophoresis. Capillary electrophoresis was performed on the 3130xl genetic analyzer using POP-7 (Thermo Fisher Scientific), and alleles were analyzed using GeneMaker v2.4.0 software (SoftGenetics, State College, PA, USA). For the other 43 SNP markers mapped to LG9, genotyping was performed using the 48.48 Dynamic Array (Fluidigm, South San Francisco, CA, USA). For the 48.48 Dynamic Array, 6.25 ng genomic DNA per sample (at a concentration of 5 ng/μL) were used for specific target amplification. The assays were performed following the protocol provided by the manufacturer. The data obtained were analyzed using Fluidigm SNP Genotyping Analysis software (ver. 4.5.1). The primer information is provided in Table S2.

Chi-square tests were performed for each locus to assess the deviation from the expected Mendelian segregation ratio. Loci showing extreme segregation distortion (*P* < 0.01) and with many missing data points (more than five individuals) were excluded from further linkage analysis. The linkage analyses were performed using the maximum likelihood mapping algorithm in JoinMap ver. 4.1 software (Kyazma, Wageningen, The Netherlands) with a cross pollination-type population (hk × hk, lm × ll, and nn × np) and two rounds of map calculation [27]. Markers were assigned to the LG9 linkage group using a logarithm of odds ratio threshold of 8.0, which was the same value as in previous reports on *C. japonica* [16, 17, 18, 26]. The maximum likelihood mapping algorithm was used to determine marker order in the linkage group. The map distance was calculated using the Kosambi mapping function [28]. Default settings were used for the recombination frequency threshold and ripple value.

### MAS of trees with *ms1*

Leaves for MAS selection were collected from breeding materials in Niigata (Tohoku breeding region), Yamagata (Tohoku breeding region), Miyagi (Tohoku breeding region), Shizuoka (Kanto breeding region), Tottori (Kansai breeding region), and Kumamoto (Kyushu breeding region) Prefectures with sample numbers of 238, 163, 30, 34, 72, and 66, respectively. In the samples from Miyagi Prefecture, Kurihara-4, a tree heterozygous for *MS1*, was included. Genomic DNA was extracted from these needles using a modified CTAB method [25, 30]. In addition, we also performed MAS selection using previously extracted DNA from 47 *C. japonica* trees in the Ishinomaki natural population of Miyagi Prefecture, where clonal analysis was performed in 2017 [29].

Based on the sequence information of CJt020762, Hasegawa et al. [30] developed two primer pairs that sandwich the two deletions, respectively. These two markers were used for MAS selection in this study. PCR amplifications were performed in 10 μL reaction volumes containing 5 ng of genomic DNA, 1× PCR Kapa2G buffer with 1.5 mM MgCl_2_, 0.2 μL of 25 mM MgCl_2_, 0.2 μL of 10 mM each dNTP mix, 0.4 μL of 5 μM forward primers labeled with dye (CJt020762_*ms1-1*_F and CJt020762_*ms1-2*_F), 0.2 μL 5 μM reverse primers (CJt020762_*ms1-1*_R and CJt020762_*ms1-2*_R), 5 ng template DNA, and 0.5 U KAPA2G Fast PCR enzyme (KAPA2G Fast PCR kit; KAPA Biosystems, Wilmington, USA). Amplification was performed on the Takara PCR Thermal Cycler (Takara) under the following conditions: initial denaturation for 3 min at 95 °C, followed by 35 cycles of denaturation for 15 s at 95 °C, annealing for 15 s at 60 °C, extension for 1 s at 72 °C, and a final extension for 1 min at 72 °C. PCR products and the DNA size marker (LIZ600; Thermo Fisher Scientific) were separated by capillary electrophoresis on the ABI 3130 Genetic Analyzer (Applied Biosystems, Tokyo, Japan). DNA fragments were detected using GeneMarker software (ver. 2.4.0; SoftGenetics).

## Results and Discussion

### Linkage maps of the *MS1* region

Of the 130 S11-S progeny produced by artificial crossing between Shindai11 and Suzu-2, 75 were male-fertile and 55 male-sterile. On the other hand, of the 138 S12-S progeny produced by artificial crossing between Shindai12 and Suzu-2, 65 were male-fertile and 73 male-sterile. The segregation ratio between male-sterile and male-fertile trees in S11-S and S12-S progenies did not deviate significantly from the expected ratios of 1:1 (X^2^ = 0.31 [*P* = 0.08] and 0.46 [*P* = 0.50], respectively). These results clearly suggested that the male-sterile gene of Shindai11 and Shindai12 was *MS1*. Based on observations using a microscope, Miura et al. [31] reported that the male-sterile phenotype of Shindai11 and Shindai12 was similar to those of Fukushima1, Fukushima2, and Shindai3, which are regulated by the *MS1* gene [6, 7]. These previous observational results obtained by microscopy are consistent with the results in this study.

The 19 and 18 markers were mapped onto the partial linkage maps of the region encompassing *MS1* for the S11-S and S12-S families, respectively (Fig. 1). There was collinearity (conserved gene order) among the two partial linkage maps. Two markers (CJt020762_*ms1-1* and reCj19250_2335) were mapped to the same position as the *MS1* locus in both maps. Of these markers, reCj19250_2335 could not be used to predict trees with *ms1* with 100% accuracy [21]. Therefore, we used CJt020762 for MAS in this study. As genome sequencing has now been conducted in *C. japonica*, the question of whether these markers are located close to each other within the genome will probably be investigated in the near future.

**Figure 1.**
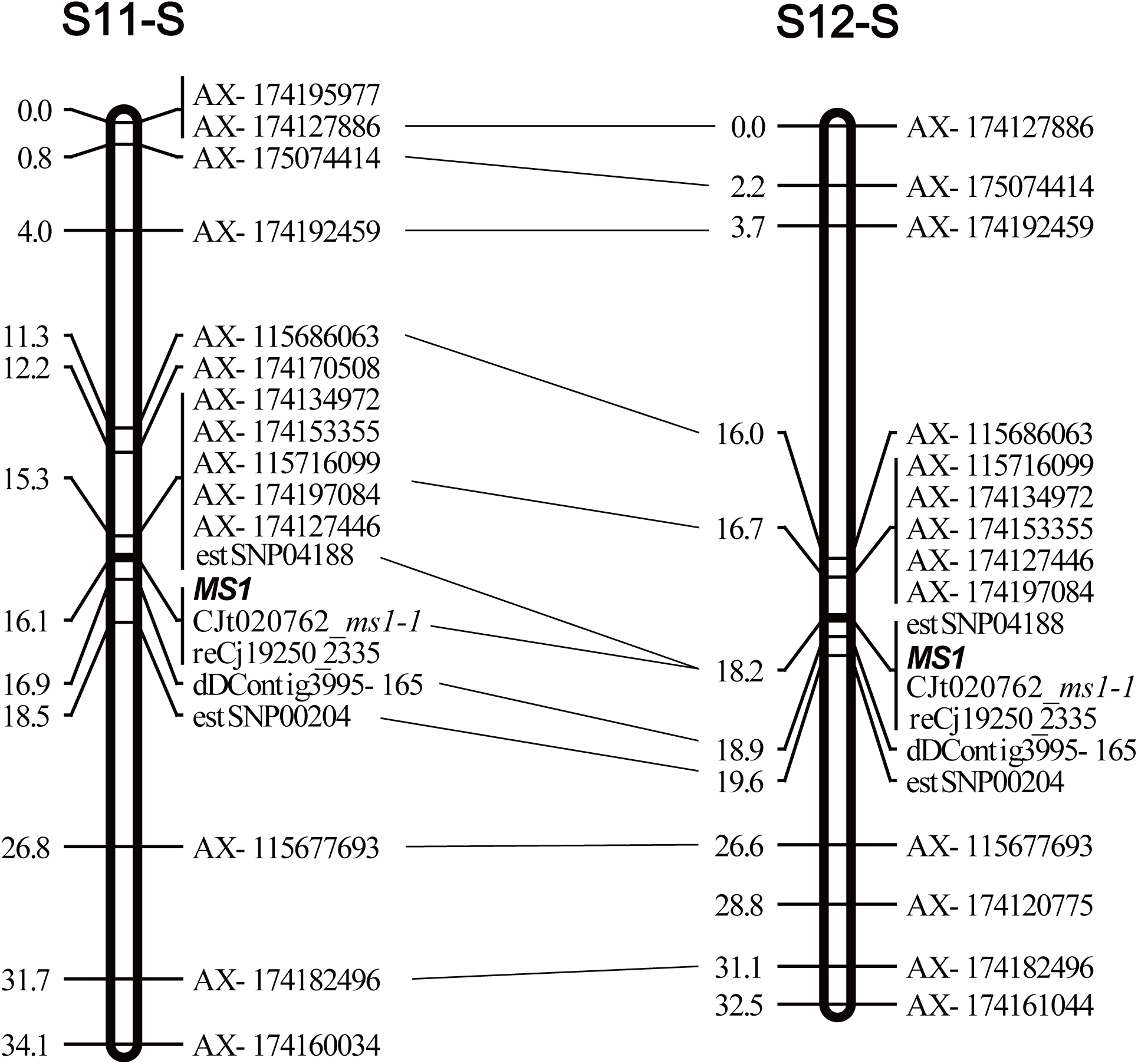
Partial linkage maps of the region encompassing *MS1* in the S11-S and S12-S *C. japonica* families.

### MAS of trees with *ms1*

In the MAS results of this study, we found that five trees in Niigata Prefecture (Kashiwazakishi-1, Setsugai Niigata-6, Setsugai Murakami-2, Setsugai Aikawa-8, and Kamikiri Niigata-55) and one tree in Yamagata Prefecture (Taisetsu Yamagata-8) had heterozygous *ms1-1*, and three trees in Miyagi Prefecture (Kurihara-4 and two trees in the natural population) had heterozygous *ms1-2*. Two male-sterile trees in Niigata Prefecture (Shindai11 and Shinadai-12) used as mother trees of mapping families had homozygous *ms1-1*. The two trees with *ms1-2* in the Ishinomaki natural forest (Ishinomaki_J284 and Ishinomaki_J278) were considered to have a parent–child relationship according to their genotypes. Hasegawa et al. [24] reported that trees with *ms1-2* may be distributed at a high frequency in this forest. Our results strongly support this suggestion. Through further selections from this natural forest, it may be possible to obtain more breeding materials for male sterility.

Because half of the offspring in the mapping family Fukushima1 (*ms1-1*/*ms1-1*) × Ooi-7 (*ms1-2*/*Ms1*) [21] showed male sterility, both of the trees with *ms1-1* and *ms1-2* can be used in a breeding program. Therefore, MAS should target both the *ms1-1* and *ms1-2* alleles. In this study, although we performed MAS in several prefectures of Japan, the prefectures in which we found trees with *ms1-1* or *ms1-2* were restricted (all trees with *ms1* were found in the Tohoku breeding region; Fig. 2). Our results suggested that there may be geographical hotspots for *ms1-1* and *ms1-2* in Niigata Prefecture and Miyagi Prefecture, respectively. Among the four breeding regions that use *C. japonica* for their artificial forests, the Tohoku breeding region has a relatively large amount of breeding materials for male sterility. However, the breeding materials for male sterility in the Kanto and Kansai breeding regions are still fewer than those in the Tohoku breeding region, and there are no breeding materials for male sterility in the Kyushu breeding region.

**Figure 2.**
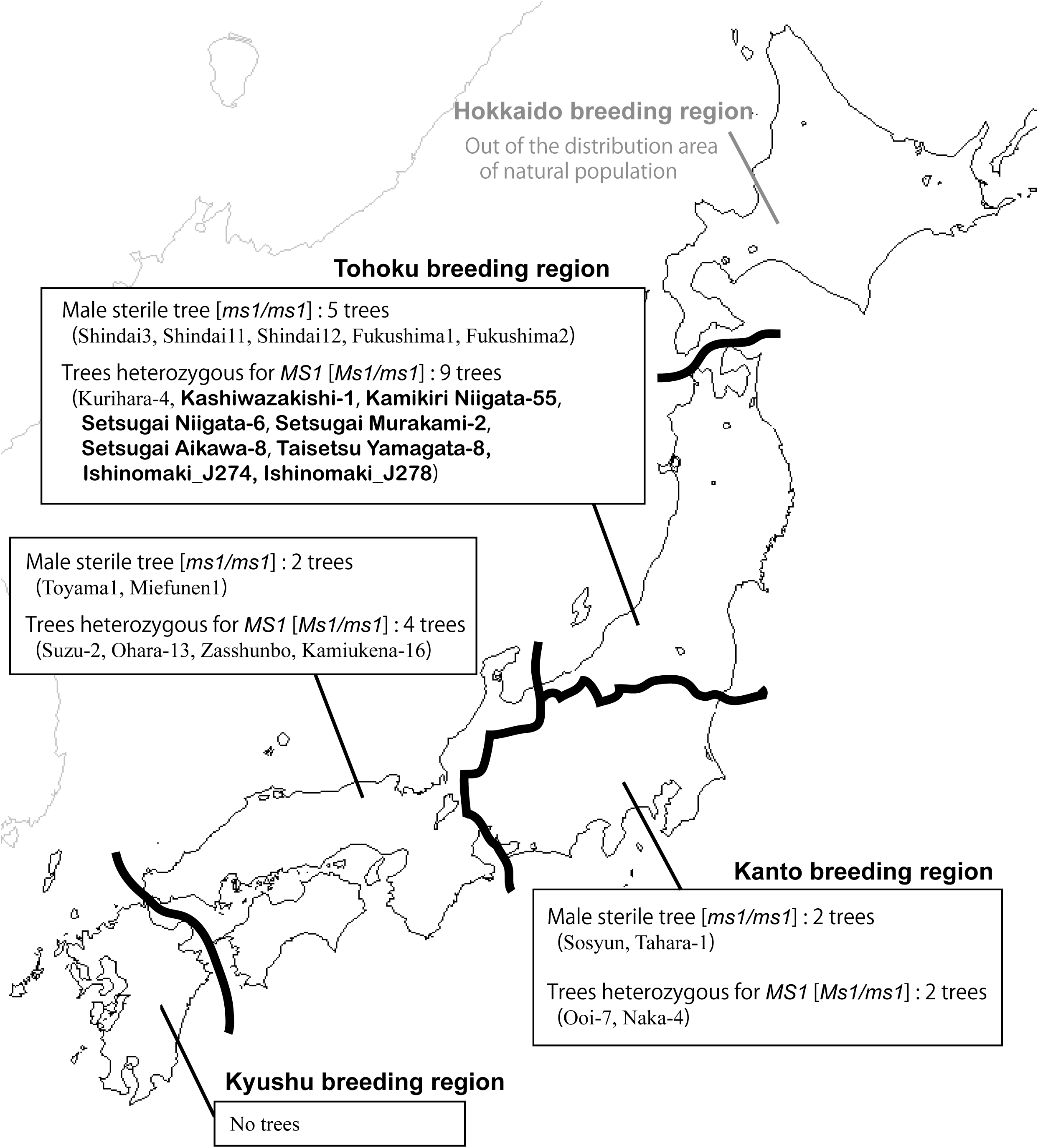
Breeding materials with *MALE STERALITY 1* of *C. japonica* in four breeding regions. The bold font shows the selected trees in this study.

It took approximately 5 years to achieve precise selection of trees heterozygous for *MS1* using a magnifying glass or a microscope (1 year to promote flowering, 1 year for seed production, and 3 years to confirm male sterility). Because MAS is effective for reducing the labor and time required for selection of trees with the male-sterile gene, the number of breeding materials should increase in the future.

## Conclusions

In this study, we performed MAS for 650 trees from six prefectures of Japan using CJt020762_*ms1-1* markers and found that five trees in Niigata Prefecture (Kashiwazakishi-1, Setsugai Niigata-6, Setsugai Murakami-2, Setsugai Aikawa-8, and Kamikiri Niigata-55) and one tree in Yamagata Prefecture (Taisetsu Yamagata-8) had heterozygous *ms1-1*, and three trees in Miyagi Prefecture (Kurihara-4 and two trees in the natural population) had heterozygous *ms1-2*. The results obtained in this study suggested that there may be geographical hotspots for the *ms1-1* and *ms1-2* alleles, respectively. Because MAS can effectively reduce the labor and time for selection of trees with the male-sterile gene, the number of breeding materials should increase in the future.

**Acknowledgements**

The authors would like to thank Y. Abe, Y. Komatsu for assistance with laboratory works. We also thank Y. Sato for artificial crossing. We also thank Y. Ito, S. Ikemoto, M. Sonoda, K. Yokoo, T. Hakamata and T. Miyashita for providing samples.

## Supplementary Materials

The following are available online at XXX, Table S1: Primer sequence of SNaPshot assay, Table S2: Primer sequence for a 48.48 Dynamic Array.

## Author Contributions

Conceptualization, Y.H., S.U., S.H. and Y.M.; Material preparation and phenotype data curation, T.T., S.H., J.I., Y.K. and Y.M.; Marker development and genotype data collection, S.U., Y.H., T.T., M.W., R.S. and Y.M.; Funding acquisition, Y.M.; Writing-original draft, Y.M.; Writing-review and editing, Y.H., S.U., M.W. and T.T. All authors have read and agreed to the published version of the manuscript.

